# Vitronectin binding affinity and cell viability effect of novel mechanotherapy drugs for neuroblastoma

**DOI:** 10.1101/2025.03.31.646300

**Authors:** Stephenie C. Alaribe, Sofia Granados-Aparici, Akolade R. Oladipupo, Isaac Vieco-Marti, Blessing E. Titilayo, Gordon Allen, Rosa Noguera

## Abstract

High-risk neuroblastoma (HR-NB) is an aggressive form of childhood cancer with a five-year survival rate of under 50%, underscoring the need for more efficacious and less toxic treatments. The glycoprotein Vitronectin (VN) has been linked to poor prognosis in patients with HR-NB, and thus inhibitors of its function represent a promising avenue for molecular mechanotherapy. The present study sought to investigate the binding affinity between the somatomedin B (SMB) domain of VN and natural compounds derived from medicinal plants. The therapeutic potential of α-amyrin (AMY), lupeol (LUP), and Olax chalcone A (Olax CHA) was tested in combination with an integrin antagonist of VN, cilengitide (CLG), using the SK-N-BE(2) HR-NB cell line as a model. Molecular docking studies indicated a potential for protein-ligand interactions for all selected compounds, of which CLG demonstrated the most favorable binding free energy (kcal/mol), followed by LUP, AMY and Olax CHA. Molecular dynamics simulations demonstrated that the SMB domain of VN initially exhibited flexibility, with alpha carbon-root mean square deviation (RMSD) stabilizing at approximately 1.8-2.1 Å. While all compounds demonstrated a dose-dependent decrease in SK-N-BE(2) cell viability, CLG exhibited higher IC50 values. Although the combination of AMY and LUP with CLG did not result in enhanced efficacy, Olax CHA exhibited a superior antiproliferative effect with higher IC50 values than AMY and LUP, and additionally showed potential synergism with CLG, suggesting a more effective therapeutic approach. This work provides valuable insights into the potential use of mechanotherapy drugs and natural products to enhance HR-NB treatment that can be expanded in future studies centered on Olax CHA.

## Introduction

Neuroblastoma (NB) is a cancer that primarily affects young children and originates in the developing neural cells of the sympathetic nervous system, commonly found in the adrenal glands, abdomen, chest, or spine (1). The disease is characterized by the uncontrolled growth of immature neural cells called neuroblasts. NB displays a wide range of clinical presentations, varying from spontaneous regression to aggressive metastatic disease. Symptoms depend on the location and extent of the tumor and may include abdominal pain, the presence of a mass or lump, bone pain, fever, fatigue, or neurological changes (2). Genetic factors, including *MYCN* gene amplification, play a significant role in NB, impacting disease aggressiveness and prognosis. Treatment approaches for high-risk NB (HR-NB) encompass a multidisciplinary strategy involving surgery, chemotherapy, radiation therapy, immunotherapy, and targeted therapies, tailored to the specific risk factors and disease stage of each patient. While advances in research and treatment have improved survival rates, challenges still exist in managing HR-NB and minimizing long-term side effects (3).

SK-N-BE(2) is a significant cell line extensively utilized in in vitro research focused on NB, other pediatric cancer treatments, and other nervous system-related cancers (4). It originated from a bone marrow metastasis of a 24-month-old child diagnosed with aggressive NB and was first documented in 1972 (5). SK-N-BE(2) cells express specific markers associated with neurofilament proteins and catecholamine production. Notably, they possess the *MYCN* amplification, a common genetic hallmark in advanced NB. This genetic alteration is frequently associated with unfavorable prognosis and aggressive disease behavior. Consequently, SK-N-BE(2) is a valuable model for investigating HR-NB biology and exploring potential therapeutic approaches.

SK-N-BE(2) cells undergo monolayer growth during their biological and culture development, with a multiplication time interval of approximately 30 hours under standard conditions (6). This characteristic makes the cell line an ideal model for investigating HR-NB, developing potential drugs, studying drug sensitivity and resistance, and making therapeutic discoveries for cancer treatment. Moreover, in recent years, SK-N-BE(2) has been extensively utilized to study various aspects of neuronal differentiation, neuronal cell signaling, and neurotoxicity assays. This expanded application showcases the versatility of SK-N-BE(2) in advancing the understanding of neurodevelopmental processes and investigating potential neurotoxic agents (7). NB cells can synthesize and secrete vitronectin (VN), including SK-N-BE(2) cells, suggesting this glycoprotein is a potential extracellular matrix (ECM) biomarker for aggressiveness in HR-NB (8). VN can bind cells through its arginyl-glycyl-aspartic acid (RGD) motif and/or domains such as its somatomedin B (SMB) domain to integrins and urokinase plasminogen activator receptor (uPAR) cell ligands (9, 10). Plasminogen activator inhibitor-1 (PAI-1) binds to the SMB domain of VN where its main function is uPA inhibition (11). Cilengitide (EMD 121974, CLG) is a mechano-drug based on the cyclic peptide cyclo (-RGDfV-), selective for ανβ3 and ανβ5 integrins. CLG and other integrin and uPAR inhibitors are still the focus of attention for the treatment of pediatric and adult cancers (12, 13).

Molecular docking and molecular dynamics simulations are important computational tools that help in predicting the binding affinity, stability, and behavior of small molecule inhibitors within their target proteins. Molecular docking sheds light on interaction modes and binding energies, while molecular dynamics offers a deeper understanding of protein-ligand dynamics over time, revealing stability and potential dissociation events (14). Some docking studies between VN (12), CLG (15) and ανβ3 receptor (15) and between growth factor domain (GFD) and SMB peptides and uPAR have been performed to identify potential binding affinities (16).

This study primarily focuses on the use of pure metabolites from medicinal plants, specifically α-amyrin (AMY), lupeol (LUP), and Olax chalcone (Olax CHA), for inhibiting the growth of SK-N-BE(2) cells in comparison with CLG, which has been previously shown to affect proliferation and adhesion dynamics of HR-NB cell lines (14, 17, 18), and also investigating their structural binding interactions with the SMB domain of VN as an alternative and potentially more effective method for inhibiting SK-N-BE(2) cell viability based on molecular mechanotherapy.

## Materials and Methods

### Docking studies

The 3D structures of the studied ligands were retrieved from PubChem in Simulation Description *Format* (SDF) and prepared using the Micromodel module of the Schrödinger suite. The resulting minimized ligands were then evaluated for their binding affinity using molecular docking studies (19). Molecular docking of the selected compounds was performed to investigate binding interactions and generate initial protein–ligand complex coordinates for subsequent molecular dynamics simulations.

The X-ray crystal structure (PDB ID 1OC0) served as the receptor. Although this is the PAI-1 domain and the SMB domain of the VN, only the SMB domain was used in these studies. The 3D structure of the target protein was prepared using Maestro’s Protein Preparation Wizard, which involved refinement steps such as removing clashes, water molecules, and unnecessary atoms, as well as adding hydrogen atoms and completing any incomplete residues. Schrödinger’s SiteMap tool was employed to identify the binding site (20). SiteMap results were instrumental in accurately locating binding sites, as they highlight hydrogen bond donors, hydrogen bond acceptors, hydrophobic regions, and hydrophilic regions. Molecular docking studies were then conducted using the extra precision (XP) Glide tool in the Schrödinger suite to ensure accurate binding pose predictions (21).

### Molecular dynamics simulations

Molecular dynamics simulations were performed using the Schrödinger LLC package. The isothermal–isobaric ensemble [NPT, an *isolated system with fixed pressure (P)*, a fixed number of atoms (N), and a fixed temperature (T)] was employed at T = 300 K and P = 1.01325 bar, with a simulation length of 100 nanoseconds (ns) and a relaxation time of 1 picosecond (ps) for all selected poses. The OPLS4 force field parameters were applied, with a cutoff radius of 9.0 angstroms (Å) for Coulomb interactions and orthorhombic periodic box boundaries set at 20 Å from protein atoms. Water molecules were described using the simple point charge (SPC) model, and a salt concentration of 0.15 M NaCl was applied using Desmond System Builder. The SPC model was used to represent water (22). P control was achieved through the Martyna-Tuckerman-Klein chain coupling scheme (coupling constant: 2.0 ps), and T control was achieved through the Nosé-Hoover chain coupling scheme (23, 24). Structural analysis of the simulation trajectory was performed using root mean square deviation (RMSD), root mean square fluctuation (RMSF), radius of gyration (Rg) and Solvent Accessible Surface Area (SASA) calculations to investigate changes occurring during the simulation (19).

### Cell culture

*MYCN*-amplified SK-N-BE(2) human NB cell lines were acquired from American Type Culture Collection (ATCC, Manassas, VA, USA) and cultured in Iscove’s Modified Dulbecco’s Medium (IMDM medium, Gibco, Life Technologies, Waltham, MA, USA) supplemented with 10% heat-inactivated fetal bovine serum, 1% insulin-transferrin-selenium and 1% penicillin/streptomycin and Plasmocin (0.2%,1/10) [InvivoGen] at 37 °C in 5% CO^2^ atmosphere.

### Cell morphology and viability assay

The natural compounds were pre-solubilized in dimethyl sulfoxide (DMSO) and test concentrations (15‒100 μM) were prepared in sterile phosphate-buffered saline (PBS). The final concentration of DMSO in the test concentrations was kept below 1%. Experimental drugs AMY, LUP, and Olax CHA were solubilized in DMSO at the desired stock concentration and CLG was dissolved in PBS. At the time of drug addition, an aliquot of the stock was thawed and diluted to twice the desired final maximum test concentration with a complete medium. Additional four, 10-fold, or ½ log serial dilutions were made and added to the appropriate wells to reach the required final drug concentrations. For drug combination studies, each of the compounds was combined with CLG (100µM) at a ratio of 1:1. Cells cultured in a latex solution at 0.05mg/ml were used as a positive control for cell death (25, 26). Cells were seeded in 8-well cell culture slides (ThermoFisher) at a density of 5×10^3^ cells in 200 μL volume. After 24 hours, cells were treated with the different concentrations of AMY, LUP, Olax CHA, and CLG for 48 h at 37 °C in 5% CO^2^ atmosphere. Phase contrast microscopy (Leica Inverted Phase Contrast microscope) was used to visualize and photograph daily changes in cell morphology and behaviors in response to drug treatment. Drug antiproliferative potential was evaluated using the alamarBlue cell viability assay protocol (ThermoScientific). Briefly, 100 μL of media was added to a 96-well clear bottom microplate (Corning, Sigma) followed by the addition of 10 μL (10% of volume in the wells) of alamarBlue reagent and kept at 37 °C in 5% CO_2_ atmosphere. Absorbance readings were recorded 4h after adding the alamarBlue reagent in a 96-well microplate reader (27) with excitation wavelength at 560 nm and emission wavelength at 590 nm.

### Drug combination studies

The inhibitory property of each combined treatment was evaluated using the combination index (CI) according to the Chou-Talalay method (28). CI values are defined as follows: CI < 1 indicates synergism; CI = 1 additivity, and CI > 1 antagonism. The dose reduction index (DRI) of AMY, LUP, Olax CHA, and CLG was calculated and defined as the measure of the dose fold decrease for each drug when synergistically combined to achieve a given effect, compared with the dose of each drug alone required to engender the same inhibition (28). Graphs and data analysis were done on GraphPad Prism (version 6.0). Results are presented as mean ± SEM of values obtained in replicate experiments.

## Results

### Molecular docking reveals interactions between selected bioactive compounds and the SMB domain of VN

Molecular docking studies provided valuable insights into the binding interactions between the investigated compounds and the SMB domain of VN (Figure 1, Tables 1–2). Binding affinities, free energies, and modes were examined, revealing that CLG exhibited the strongest overall binding affinity, with a Glide score of 5.176 kcal/mol and binding free energy (ΔGBind) of −31.89 kcal/mol (Table 1). This robust binding was attributed to strong electrostatic interactions (ΔGCoulomb = −30.36 kcal/mol) and favorable Van der Waals interactions (ΔGVdW = −35.86 kcal/mol) [Table 1]. CLG forms multiple hydrogen bonds with the amino acid residues THR161 (1.98 Å), ALA156 (2.03 Å), ALA318 (2.04 Å), PHE299 (1.84 Å), and GLU291 (1.99 Å) [Figure 1A and Table 2]. The remaining compounds showed weaker binding compared to CLG (Table 1). No hydrogen bonds were observed for AMY (Figure 1C). LUP also showed hydrogen bonds with the amino acid residues PHE299 (1.75 Å) [Figure 1E and Table 2] and Olax CHA with GLU291 (1.61 Å) and GLN301 (2.04 Å) [Figure 1G and Table 2]. The interaction profiles of the compounds during the simulation also elucidated their binding behavior (Figure 1). For CLG, amino acid residue GLU291 interacted with the hydroxyl group via hydrogen bonding for over 54% of the simulation time, indicating a moderately stable interaction (Figure 1B). In contrast, AMY interacted with the hydroxyl group through ALA156 (37%) and GLY155 (34%), indicating moderate interaction stability (Figure 1D). For LUP, no interactions with protein residues exceeded 10% of the simulation time, suggesting that this compound does not establish strong or stable interactions with the SMB domain of VN (Figure 1F). With respect to Olax CHA, it was observed that amino acid residue GLY_482_ plays a significant role in stabilizing it through its carbonyl oxygen, interacting over 49% of the simulation time (Figure 1H). This relatively stable interaction suggests that the carbonyl oxygen acts as a critical binding site, forming hydrogen bond interactions with GLY482 and allowing proper orientation within the receptor’s active site. Additionally, one hydroxyl group of Olax CHA formed a hydrogen bond with residue GLU_481_. As a negatively charged residue, GLU481 is well-suited to interact with the hydroxyl proton, reinforcing the binding stability and possibly orienting it favorably within the binding pocket. This interaction suggests that the hydroxyl group is accessible and might be functional in binding and activity.

**Figure 1.**
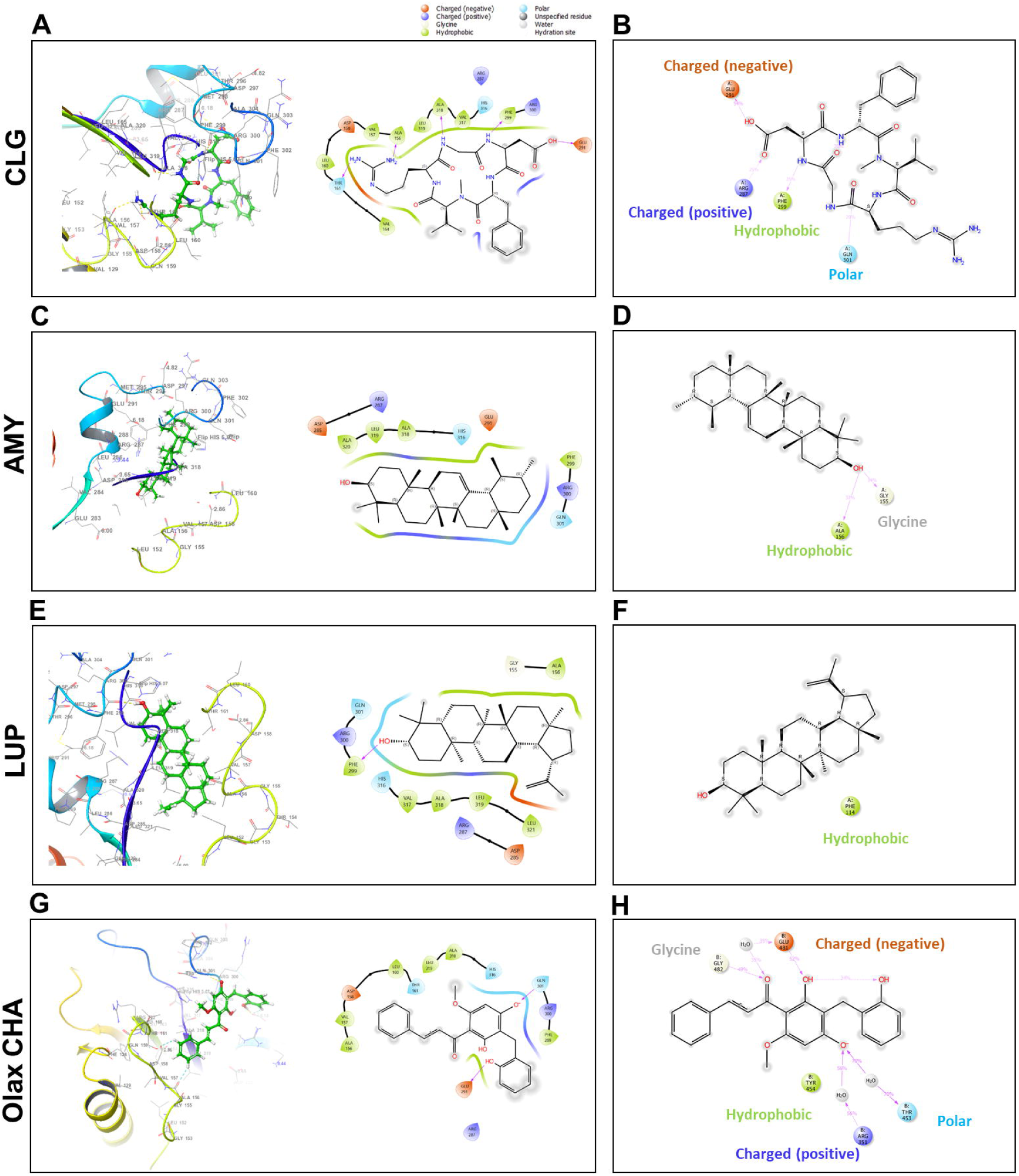
Molecular docking interactions (right) and 3D binding models (left) between the selected compounds and the SMB domain of VN. CLG (A and B), AMY (C and D), LUP (E and F) and Olax CHA (G and H). CLG: Cilengitide, AMY: α-Amyrin, LUP: Lupeol, Olax CHA: Olax chalcone A.

**Table 1:** Docking scores and binding free energy components for ligand–protein interactions.

| Entry | G-Score | $\Delta G_{\text{Bind}}$ | $\Delta G_{\text{Coulou}}$ | $\Delta G_{\text{Covalent}}$ | $\Delta G_{\text{Lipho}}$ | $\Delta G_{\text{Solv\_GB}}$ | $\Delta G_{\text{VdW}}$ |
| --- | --- | --- | --- | --- | --- | --- | --- |
| Cilengitide | -5.176 | -31.89 | -30.36 | 5.11 | -3.44 | 35.64 | -35.86 |
| α-Amyrin | -1.062 | -17.90 | -0.56 | 0.17 | -9.83 | 19.08 | -26.76 |
| Lupeol | -2.227 | -21.36 | -7.35 | 2.15 | -9.02 | 19.78 | -26.36 |
| Olox<br>chalcone A | -3.073 | -14.25 | -36.97 | 0.77 | -8.19 | 60.98 | 60.98 |

$\Delta G_{\text{Bind}}$ : Binding free energy, $\Delta G_{\text{Coulou}}$ : Coulombic energy (electrostatic interactions between ligand and protein), $\Delta G_{\text{Covalent}}$ : Covalent bonding energy, $\Delta G_{\text{Lipho}}$ : Lipophilic energy (contribution from hydrophobic/lipophilic interactions), $\Delta G_{\text{Solv\_GB}}$ : Solvation energy (generalized Born solvation model, energy required to desolvate ligand and protein) and $\Delta G_{\text{VdW}}$ : Van der Waals energy.

**Table 2:** Hydrogen bonding and π-π stacking interactions of selected compounds with vitronectin.

| ENTRY | Hydrogen bond (Å) |
| --- | --- |
| Cilengitide | THR161(1.98), ALA156(2.03),<br>ALA318(2.04), PHE299(1.84),<br>GLU291(1.99) |
| α-Amyrin | - |
| Lupeol | PHE299(1.75) |
| Olox chalcone A | GLU291 (1.61), GLN301(2.04) |

An intramolecular interaction was observed between two adjacent hydroxyl groups within Olax CHA, which formed hydrogen bonds about 34% of the simulation time. These intramolecular hydrogen bonds indicate conformational flexibility within Olax CHA, allowing it to adapt to the binding environment. This flexibility may increase its potential for binding by enabling multiple favorable interactions with different receptor regions. Intramolecular hydrogen bonding could also influence the compound’s stability, solubility, and reactivity, potentially shielding parts of the molecule from solvent interactions.

Although different binding energies were found for the different compounds, most of the compounds form one or more hydrogen bonds with different amino acid residues in the active site of the SMB domain of VN, contributing significantly to their binding affinity (Table 2). However, these findings suggest varying degrees of interaction stability across the compounds, with CLG exhibiting the most stable protein-ligand interaction.

### Molecular dynamics simulations show SMB domain flexibility and favorable binding dynamics with CLG

The RMSD values for the Cα atoms in all complexes relative to their initial structure (24) were calculated to investigate the impact of compounds on the conformational stability of the SMB domain of VN during the simulations. The results, plotted against simulation time (Figure 2), showcase the RMSD profiles of the studied compounds bound to the SMB domain of VN. To measure the statistical stability of the system, five independent simulations were performed for each pose, starting from the same energy-minimized structure (29).

**Figure 2.**
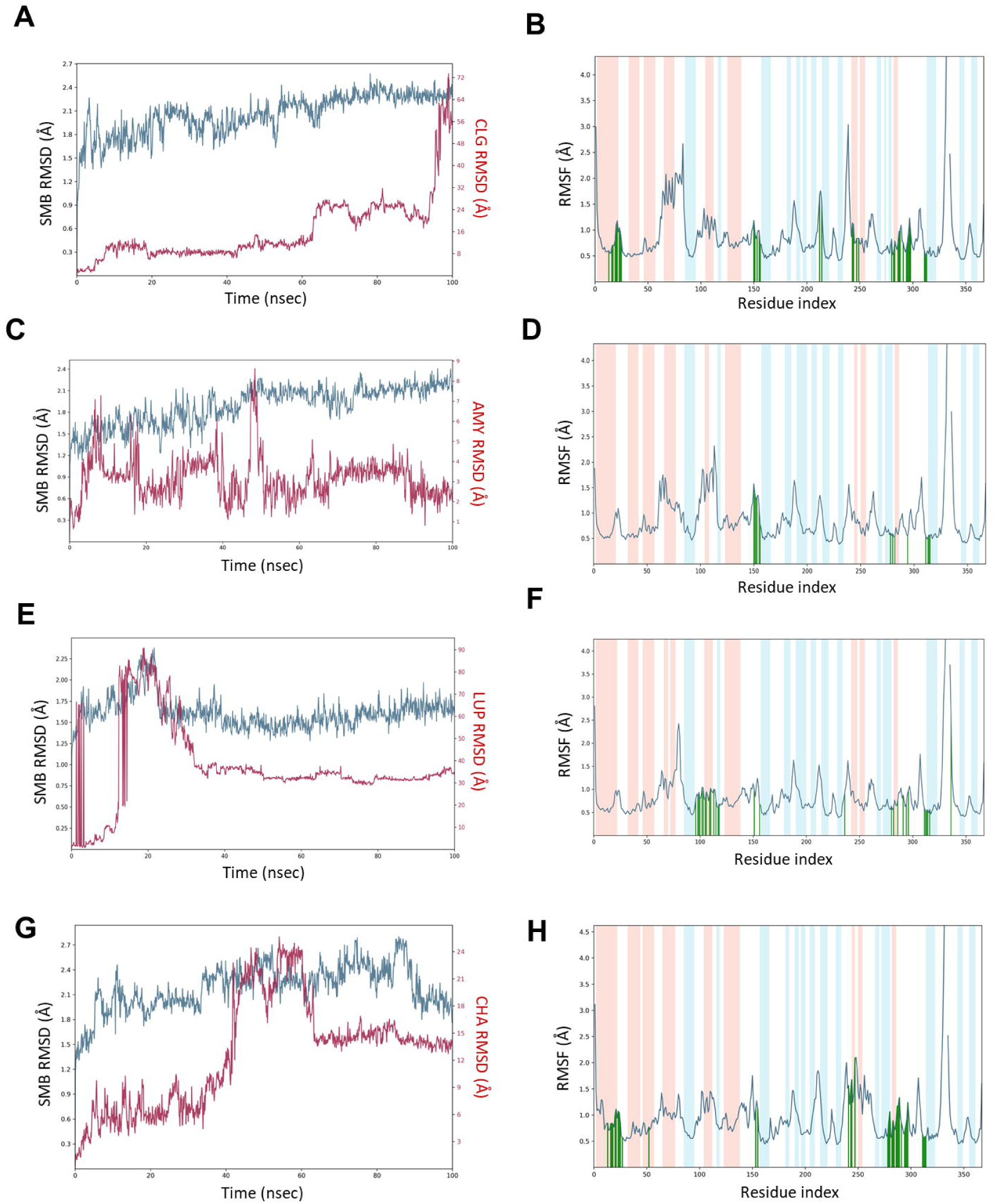
RMSD (A,C,E,G) and RMSF (B,D,F,H) profiles of the c-α and ligand fit for the following ligands with the SMB domain of VN. RMSF profiles show protein residues that interact with the ligand with green-colored vertical bars. CLG (A and B), AMY (C and D), LUP (E and F) and Olax CHA (G and H). CLG: Cilengitide, AMY: α-Amyrin, LUP: Lupeol, CHA: Olax chalcone A.

The SMB domain structure exhibited initial flexibility, as evident from the rapid rise in C-α RMSD from 0.9 Å to 1.5 Å within the first 2 ns. This was followed by a relatively stable phase, characterized by minor fluctuations in SMB RMSD (1.8 to 2.1 Å) between 2 ns and 54 ns. The SMB domain further stabilized between 54 ns and 64 ns, with reduced RMSD fluctuations at around 1.8 Å. CLG binding around 92 ns appeared to influence protein dynamics, as evidenced by increased RMSD fluctuations up to 2.1 Å. In contrast, the CLG RMSD profile showed significant fluctuations, peaking at 4 Å, 10 Å, and eventually, 24–32 Å, suggesting a possible dissociation from the SMB domain at around 90 ns, marked by a sharp RMSD increase up to 70 Å. Notably, the SMB domain and ligand CLG only interacted at around 92 ns, highlighting a binding event where the ligand interaction briefly influences SMB domain stability before dissociating (Figure 2A). Regarding AMY, the SMB RMSD reached stability at 2.4 Å, while the AMY’s RMSD achieved at around 9 Å (Figure 2C). In the case of LUP, the SMB RMSD profile stabilized at 2.25 Å, while the LUP RMSD peaked at 90 Å, after which both achieved stability at around 40 ns, remaining stable for the rest of the simulation (Figure 2E). Interaction with Olax CHA showed minor fluctuations, with the SMB RMSD achieving stability at around 4 Å and the Olax CHA RMSD at around 5.6 Å, suggesting a relatively stable contact with minimal sporadic oscillations (Figure 2G). These observations suggest a dynamic interplay between the SMB domain and the selected ligands, with binding influencing the SMB domain and ultimately leading to ligand dissociation.

Protein dynamics were also analyzed by examining RMSF, which are calculated relative to the average conformation from molecular dynamics simulations, serving as an indicator of flexibility differences between residues (30). The RMSF profiles of the C-α atoms of the analyzed compounds yielded important insights into their structural flexibility and stability (Figure 2). For CLG, the RMSF remained below 1.3 Å within the first 50 residues, with a slight increase to 1.5–2.0 Å between residues 50 and 90, suggesting a relatively stable structure early on, followed by moderate flexibility (Figure 2B). AMY showed moderate flexibility (0.5–2.0 Å), indicating subtle differences in conformational behavior and interaction capabilities (Figure 2D). LUP demonstrated a more rigid structure, with fluctuations primarily between 0.5 and 1.0 Å, but an occasional increase to 2.5 Å suggested regions of higher flexibility (Figure 2F). This combination of rigidity and occasional flexibility may be critical for its interaction with biological targets. Olax CHA induces notable flexibility in specific regions of the domain, with RMSF peaks of up to 4 Å around residues 50, 150, and 220, suggesting it promotes conformational adjustments that may be essential for binding.

To assess the structural stability and degree of compactness of the SMB domain–CLG complex, the relative radius of gyration (rGyr) and global solvent accessible surface area (SASA) were calculated (31). The rGyr, which provides data on the kinetics and thermodynamics of protein folding, serves as a key indicator of protein compactness and overall stability of biomolecular structures (32). Figure 3A illustrates the ligand properties of CLG, highlighting several key features such as rGyr and SASA. An in-depth analysis of these properties revealed that the CLG RMSD fluctuated between 1 and 3 Å and displayed oscillations in these hydrogen bonds between 0 and 3 Å, stabilizing at around 1 Å for most of the simulation time. This suggests a dynamic hydrogen-bonding pattern in CLG that contributes to its overall stability. Regarding the degree of compactness, the CLG SASA values ranged from 450 to 900 Å², stabilizing at approximately 450 Å² between 20 ns and 40 ns of the simulation, indicating rather low compactness, particularly over the latter stages of the simulation.

**Figure 3.**
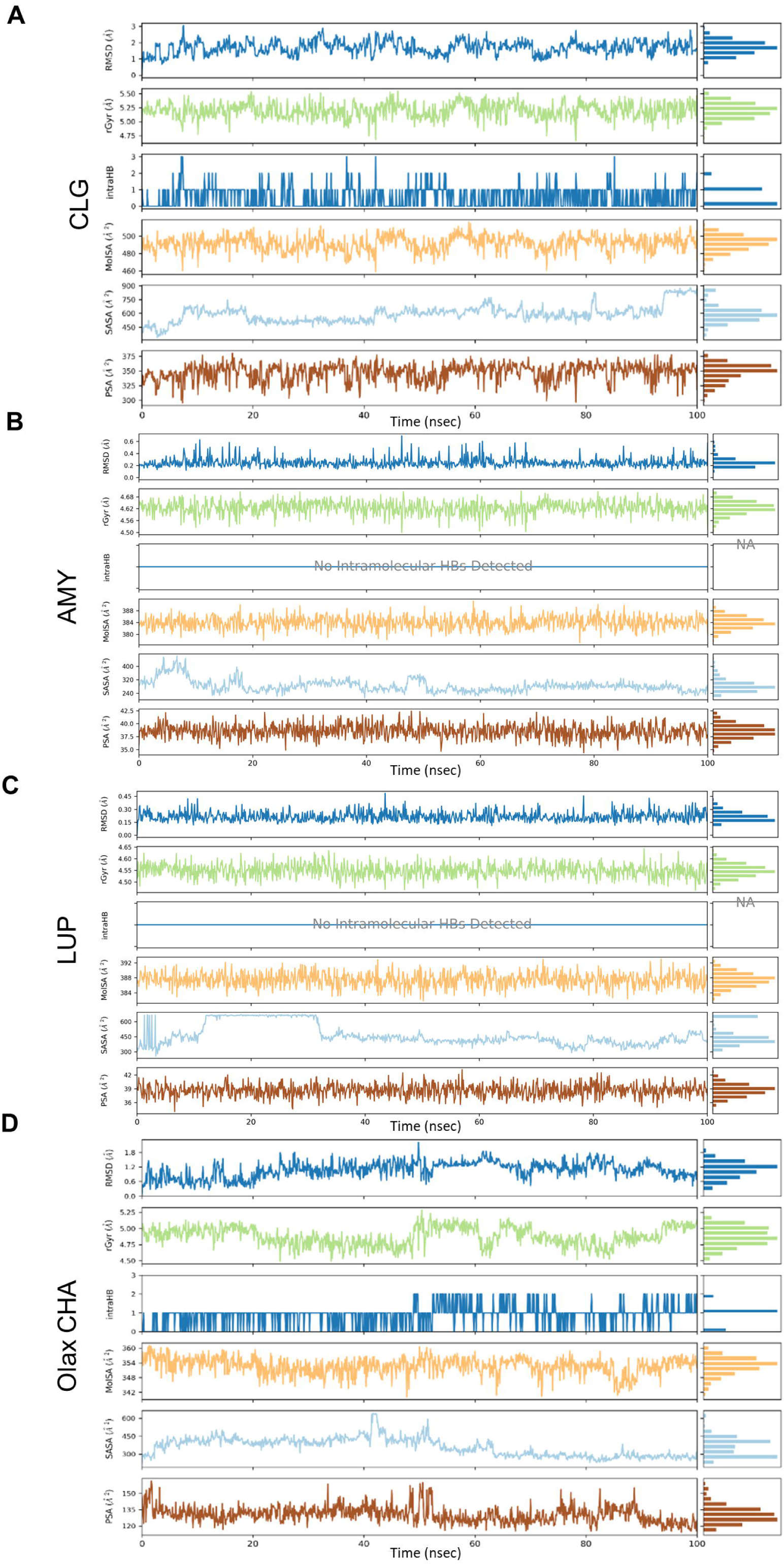
Diagram comparing ligand properties of CLG (A), AMY (B), LUP (C) and Olax CHA (D) featuring ligand RMSD, radius of gyration (rGyr), intramolecular hydrogen bonds (intraHB), molecular surface area (MolSA), solvent accessible surface area (SASA), and polar surface area (PSA). CLG: Cilengitide, AMY: Amyrin; LUP: Lupeol; Olax CHA: Olax chalcone A.

In rGyr measurements of ligand compactness, AMY exhibited values ranging from 4.56 to 4.68 Å while LUP maintained a slightly lower and consistent range between 4.50 and 4.60 Å (Figures 3B and 3C, respectively). Neither ligand formed intramolecular hydrogen bonds during the simulation, consistent with their predominantly hydrophobic and sterically bulky structures. Molecular surface area (MolSA) values for AMY ranged from 380 to 388 Å^²^, while LUP displayed slightly higher values of 384 to 392 Å^²^, reflecting minor differences in molecular geometry. The AMY SASA ranged from 240 to 400 Å², suggesting tighter packing and reduced solvent exposure. SASA values for LUP were more dynamic, fluctuating between 300 and 600 Å^²^, with a brief stabilization above 600 Å^²^ between 15 ns and 37 ns. The PSA values of both ligands remained relatively narrow, with AMY ranging between 35 and 42.2 Å^²^ and LUP between 36 and 42 Å^²^, indicating minimal polar character and consistent interaction profiles. Overall, LUP exhibited slightly lower solvent exposure dynamics, possibly due to transient repositioning or a more flexible binding environment. In contrast, the tighter packing observed with AMY suggests stronger hydrophobic interactions and a more rigid interaction profile. These findings highlight the distinct interaction patterns of LUP and AMY, helping unravel their potential binding mechanisms and functional roles within the simulated system.

Conversely, for Olax CHA, the protein RMSD started below 1.5 Å and spiked to about 2.4 Å in the first 15 ns of the run time, suggesting structural relaxation or conformational adjustments of the protein upon ligand binding or simulation initialization. After this, the protein stabilized momentarily between 17–33 ns at around 1.8 Å, and then fluctuated for the remainder of the simulation, as shown in Figure 3D. This simulation broadly explains the interaction dynamics between the protein and ligand over the 100 ns trajectory, our observations indicating that the protein maintains an overall stable structure, although localized flexibility likely occurs due to environmental factors such as solvent effects or ligand interactions. In contrast, the ligand RMSD demonstrated erratic behavior, with transient binding to the protein’s active site observed only between 40 and 60 ns, where it fluctuated between 18 Å and 24 Å. This suggests that the ligand engages the active site weakly and intermittently, potentially exploring various binding poses or undergoing displacement during the simulation.

The RMSF profile further supports the stability of the protein, with residue fluctuations remaining below 2.0 Å for the first 300 residues. Notably, key residues LYS288 and GLU291 emerged as critical interaction points. Hydrogen bonding between the ligand’s carbonyl oxygen and LYS288, as well as interactions involving its hydroxyl groups with GLU291, highlight their roles in ligand anchoring. However, the transient nature of these interactions, observed at 46% and 39% of the simulation time, respectively, reflects limited ligand retention. Additionally, intramolecular hydrogen bonding within the ligand may compete with its ability to form stable interactions with the protein.

Further analysis of ligand properties revealed intrinsic flexibility, with its RMSD fluctuating between 0.6 Å and 1.8 Å. The rGyr variation from 4.50 to 5.25 Å indicates dynamic changes in ligand compactness during the simulation. Moderate solvent exposure, as evidenced by molecular surface area (MolSA: 342–360 Å^²^), SASA: 300– 600 Å^²^, and polar surface area (PASA: 120–150 Å^²^), reflects the dynamic nature of ligand-protein interactions.

### CLG, AMY, LUP, and CHA treatment show a dose-dependent antiproliferative effect in SK-N-BE(2) cells

Phase contrast micrographs were acquired at 24h and 48h after treatment with 30µM and 100µM of CLG, AMY, LUP, and Olax CHA to study their effect on cell morphology (Figure 4A–B). The control conditions (no treatment) showed the expected morphology and growth rate of the SK-N-BE(2) cells, which exhibited a range of morphologies, including processes of varying length and an epithelioid-like arrangement. Cell aggregation was observed, resulting in the formation of clumps that exhibited a tendency to float. A mean multiplication time interval of approximately 30 hours was recorded. In contrast, treatment with a solution incubated with latex resulted in minimal proliferation rate of 24–48 hours. Treatment with 100 µM CLG showed similar results to those recently reported (18): rapid cell aggregation as a consequence of induced cell detachment with a long multiplication time interval. Treatment with AMY, LUP, or Olax CHA did not result in any notable differences in morphology or growth rate for controls at any of the time points or concentrations (Figure 4A). However, when each treatment was combined with 100µM of CLG, cells showed a slight trend towards clustering together rather than covering the spaces available, similar to the effect observed under exposure to CLG alone (Figure 4B).

**Figure 4.**
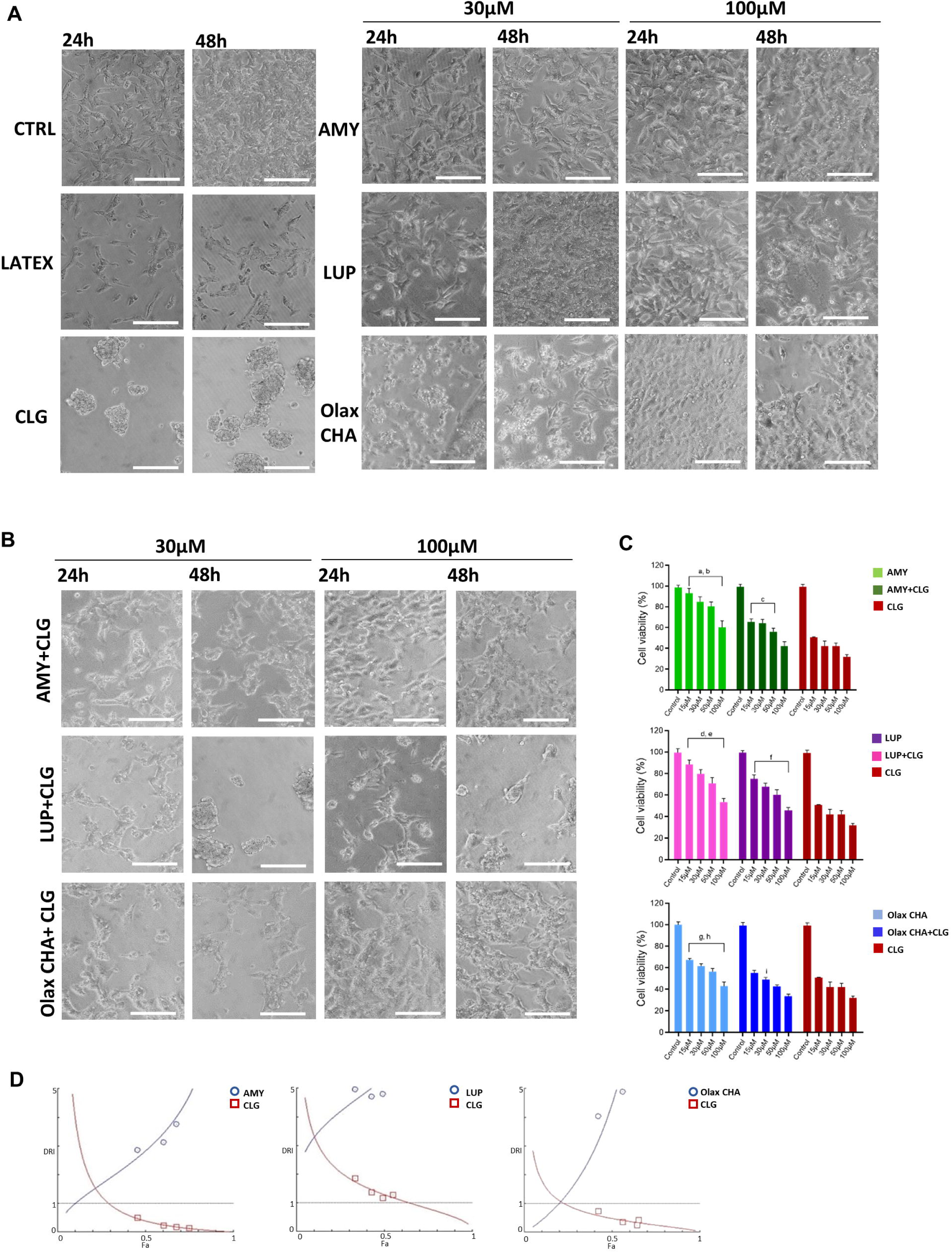
Morphological, cell viability studies, and combination index studies. A) Phase contrast microscopy micrographs at 24h and 48h of SK-N-BE (2) cell treatment with 30uM and 100um of AMY, LUP and Olax CHA. Latex was used as a control for cell death and CLG was used as the control for cell detachment. B) Phase contrast microscopy micrographs at 24h and 48h of cell treatment with 30µM and 100µm of AMY, LUP, and Olax CHA in combination with 100µM of CLG. C) Cell viability effects of the combination of 100µM of CLG with 30µM and 100µm of AMY, LUP, and Olax CHA. Scale bar is 100µm. D) Graphical representation of dose reduction index (DRI) analysis of AMY, LUP and CHA in combination with CLG. Fa: Fraction of cells affected. AMY: α-Amyrin, LUP: Lupeol, Olax CHA: Olax chalcone A and CLG: Cilengitide.

To better assess cell viability following treatment and its impact on cell proliferation or metabolic activity, we employed alamarBlue reagent. This tests the ability of viable cells to reduce resazurin (a non-toxic, cell-permeable compound) to resorufin, which can be quantified by measuring absorbance. Following a 48-hour incubation period, we assessed the viability of cells treated with varying concentrations of CLG, AMY, LUP, and Olax CHA, also obtaining the corresponding IC50 values (Figure 4C and Table 3). The results demonstrated a notable decline in cell viability following treatment with CLG. At the lowest concentration, there was a mean reduction to 50.86% viability, while at 100 µM, viability fell to 31.87%. This trend was found to correlate with IC50 values, which were closer to the lowest concentration tested (15.25 ± 1.9). The viability of cells treated with AMY showed only a slight decrease with increasing concentrations, ranging from 93.19% to 60.19%. IC50 values were beyond the range of concentrations employed (125.45 ± 8.9). A comparable pattern was evident for LUP, with concentrations spanning from 88.33–53.76% mean cell viability and elevated IC50 values (103.36 ± 1.9). Olax CHA treatment had a more pronounced impact on cell viability than AMY and LUP, showing 67.4% to 42.9% mean cell viability and lower IC50 values (74.23 ± 4.7).

**Table 3.** IC50 values of CLG, AMY, LUP, and Olax CHA and their combinations on SK-N-BE(2) cell line.

| Drug molecule(s) | IC<sub>50</sub> (μM) |
| --- | --- |
| CLG | 15.25 ± 1.9 |
| AMY | 125.45 ± 8.9 |
| AMY + CLG | 67.92 ± 2.3 |
| LUP | 103.36 ± 1.9 |
| LUP + CLG | 84.46 ± 1.3 |
| CHA | 74.23 ± 4.7 |
| CHA + CLG | 26.47 ± 1.4 |

AMY: α-amyrin, LUP: lupeol, CHA: chalcone, CLG: cilengitide.

When AMY, LUP, and Olax CHA were combined with 100µM of CLG, mean cell viability and IC50 values were obtained with a CI and DRI to explore the possibility of a synergistic interaction between compounds. All three products showed a decrease in cell viability and IC50 values (Figure 4C, Figure 4D and Table 3). However, while AMY and LUP exhibited CI and DRI values above 1 (indicative of an absence of synergy between compounds), CGL DRI values were below 1, indicating an antagonistic or additive effect rather than a synergistic interaction (Table 4, 5). In contrast, the CI values for Olax CHA showed a progressive decrease as the concentration increased, falling from 1.06 at 30 µM to 0.74 at 100 µM (Table 6). This indicates a shift from additivity towards synergism between Olax CHA and CLG at these concentrations. Additionally, the DRI values for both Olax CHA and CLG were greater than 1 (Table 6), reflecting a synergistic interaction where the combined effect is greater than the sum of the individual effects.

**Table 4.** Results of synergy study of cilengitide (CLG) and α-amyrin (AMY) against SK-N-BE(2) cell line.

| **Dose (μM)** | **CI** | **DRI AMY** | **DRI CLG** |
| --- | --- | --- | --- |
| 15 | 2.39 | 11.57 | 0.43 |
| 30 | 4.34 | 5.99 | 0.24 |
| 50 | 3.10 | 4.87 | 0.34 |
| 100 | 1.62 | 4.03 | 0.73 |

CI: combination index, DRI: dose reduction index

**Table 5.**
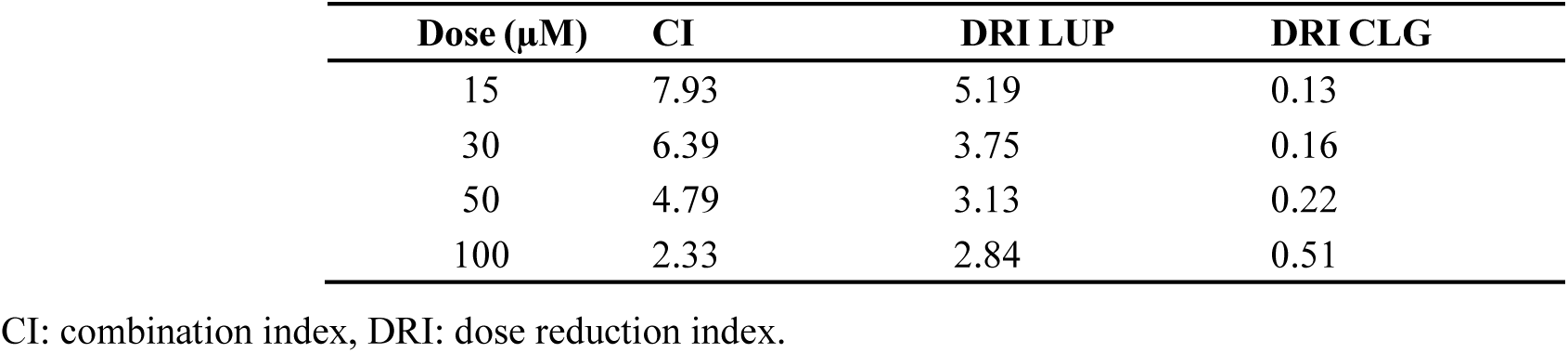
Results of synergy study of cilengitide (CLG) and lupeol (LUP) against SK-N-BE(2) cell line.

**Table 6.**
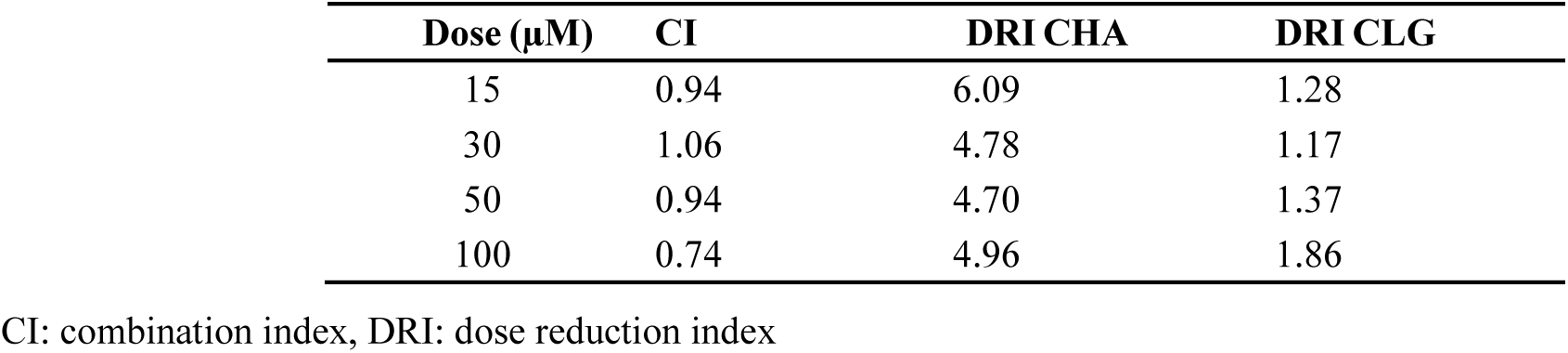
Results of synergy study of cilengitide (CLG) and Olax chalcone A (Olax CHA) against SK-N-BE(2) cell line.

| **Dose (μM)** | **CI** | **DRI CHA** | **DRI CLG** |
| --- | --- | --- | --- |
| 15 | 0.94 | 6.09 | 1.28 |
| 30 | 1.06 | 4.78 | 1.17 |
| 50 | 0.94 | 4.70 | 1.37 |
| 100 | 0.74 | 4.96 | 1.86 |

CI: combination index, DRI: dose reduction index

## Discussion

Despite representing only about 6% of childhood cancers, and the advancements in treatment options, HR-NB continues to pose a significant threat, accounting for approximately 15% of all pediatric cancer deaths (39). The five-year survival rate for HR-NB remains below 50%, highlighting a critical need for more effective and less toxic therapies (40). To enhance the efficacy of chemotherapeutic approaches in SK-N-BE(2) treatment, targeted therapies have been utilized. These therapies aim to inhibit specific signaling pathways, such as the *MYCN* gene pathway, which contribute to HR-NB development (41). However, the effectiveness of targeted therapy can be limited by the genetic heterogeneity observed among different NB tumors. This heterogeneity involves genetic alterations and variations that affect the presence or activity of targetable molecules. Consequently, the response to targeted therapy may vary among patients based on their specific genetic profiles. While chemotherapeutic agents remain an important part of HR-NB treatment, their limitations highlight the need for alternative approaches, such as combining them with the natural compounds tested in this study, which can offer enhanced selectivity and diminished toxicity (42).

CLG is a potent and selective anti-angiogenic small molecule targeting the αvβ3, αvβ5, and α5β1 integrins (50). Various studies have demonstrated its potential as an anti-cancer and anti-angiogenesis drug (51, 52). However, a phase III clinical trial (CENTRIC EORTC 26071-22072) combining CLG with temozolomide in glioblastoma patients failed to show improvement in overall survival (53). This limited clinical efficacy was attributed to the drug’s rapid clearance and short half-life, a limitation that can be effectively addressed through nanoencapsulation (14). In a recent study, etoposide showed strong synergy when co-administered with CLG in both free and nanoencapsulated form against SK-N-BE(2) and NBL-S NB cells (14). This suggests that CLG-based combination therapies offer a promising approach to reducing reliance on conventional chemotherapy for HR-NB patients. Furthermore, nanoencapsulation holds potential for improving the pharmacokinetics and bioavailability of CLG and potentially other combination agents. This study aimed to explore the potential benefits and advantages of using natural products such as AMY, LUP, and Olax CHA plus CLG as combination measures for inhibiting proliferation and/or viability of the human NB SK-N-BE(2) cell line.

Molecular docking experiments revealed interactions between selected bioactive compounds and the SMB domain of VN, illustrating the potential of these compounds to modulate the VN–uPAR–PAI-1 interactions, which are critical in regulating fibrinolysis, angiogenesis, and cancer metastasis (43, 44). Ouyang et al. (2021) extensively reviewed the in vitro and in vivo anticancer activities of CHA and its derivatives, which include cell cycle disruption, autophagy regulation, and apoptosis induction (45). Our study has demonstrated that CLG and CHA exhibited the highest binding free energy. CLG’s strong binding was enhanced by hydrogen bonds with THR161, ALA156, and GLU291. The docking results correlated with in vitro IC50 data, with CLG showing the lowest IC50. Although CHA did not rank highly in docking studies, its combination with CLG resulted in lower IC50 values, showing synergistic interactions not predicted by docking simulations alone, given the well-documented anticancer potential of CHA and its derivatives. Therefore, the improved efficacy of the CHA–CLG combination suggests a synergy that may amplify their individual anticancer effects, potentially offering a more potent anticancer activity than either compound used in isolation.

The RMSD is a widely employed metric for quantifying differences in dynamics, particularly in the context of molecular dynamics trajectory analyses (46). We report here that molecular dynamics simulations revealed initial SMB domain flexibility. SMB RMSD profiles in combination with CLG and Olax CHA showed evident stability. While CLG simulation showed a significant RMSD increase at 92 ns, indicating potential ligand separation and suggesting decreased binding stability over time, the interaction with Olax CHA showed minimal fluctuations in the SMD RMSD, indicating a stable contact. The observed synergy between Olax CHA and CLG, as evidenced by CI and DRI values, could be attributed to conformational changes or transitory binding events not captured in the initial docking studies, and the potential need for molecular modification of CLG or combination therapy.

These findings contribute to the growing body of research exploring natural products as potential therapeutic agents for HR-NB (47, 48). The results further substantiate previous reports suggesting that Olax CHA exhibits more effective inhibitory activity against selected cancer targets than AMY and LUP (49, 50). While AMY and LUP did not exhibit synergy with CLG, Olax CHA demonstrated promising results. The moderate independent activity of AMY and LUP observed herein concurs with previous studies investigating their effects on various cancer cell lines (51–53). However, the lack of synergy with CLG in this study suggests further investigation is necessary to optimize their potential application in HR-NB therapy.

The superior cell viability effect of Olax CHA compared to AMY and LUP is noteworthy, and its observed shift towards synergy when combined with CLG is particularly encouraging. This finding aligns with an emerging body of research highlighting the potential of botanical extracts with Olax CHA for synergistic anticancer effects (48, 49). Olax CHA has been reported to exert a strong synergistic effect on tumor cells, including pluritherapy-resistant tumors (54). These findings provide preliminary evidence for Olax CHA’s utility as a candidate for enhancing CLG effectiveness against SK-N-BE(2) cell viability and/or proliferation.

Future research should focus on the synthesis of Olax CHA and derivatives, which can be used to explore the underlying mechanisms by which it exerts its antiproliferative effects and potentiates CLG activity through in vivo studies. Additionally, in vivo studies using relevant animal models are crucial to assess the efficacy and safety of Olax CHA, alone and in combination with CLG. Investigating its potential for dose reduction or improved therapeutic outcomes in combination with conventional therapies is also of high interest. By building upon these findings, subsequent studies could move towards developing Olax CHA-based therapeutic strategies for improved clinical outcomes in HR-NB patients.

## Conclusion

This study investigated the potential of natural compounds and other mechanotherapy drugs such as CLG as potential combination therapies for HR-NB. While AMY and LUP demonstrated moderate antiproliferative activity against the SK-N-BE(2) NB cell line, they did not exhibit synergistic effects when combined with the anti-cancer agent CLG. In contrast, Olax CHA displayed a superior independent antiproliferative effect compared to the other two natural products. More importantly, the combination of Olax CHA and CLG showed a promising shift towards synergy, suggesting a potentially more effective therapeutic approach. Further studies are warranted to elucidate the mechanisms underlying Olax CHA antiproliferative activity and its synergistic interaction with CLG. In vivo studies using relevant animal models are also necessary to evaluate the efficacy and safety of CHA, both as a single agent and in combination with CLG. Future research expanding from studies such as ours can further explore the potential of Olax CHA-based combination therapies as a novel mechanotherapy approach to improve clinical outcomes for HR-NB patients.

## Acknowledgments

The authors are grateful to the Center for High Performance Computing, CSIR Campus, 15 Lower Hope Street, Rosebank, Cape Town, South Africa. We also gratefully acknowledge support from ISCIII (FIS) and FEDER (European Regional Development Fund), grant number PI20/01107, FNB (Fundación Neuroblastoma), grant number PRV/00166, and CIBERONC (contract CB16/12/00484), Ministry of Science, Innovation and Universities of Spain (FPU20/05344). The authors thank Kathryn Davies for English corrections.

